# Identifying the genes impacted by cell proliferation in proteomics and transcriptomics studies

**DOI:** 10.1101/2022.03.15.483931

**Authors:** Marie Locard-Paulet, Oana Palasca, Lars Juhl Jensen

**Affiliations:** Novo Nordisk Foundation Center for Protein Research, University of Copenhagen, Denmark

## Abstract

Today, hypothesis-free high-throughput profiling allows relative quantification of thousands of proteins or transcripts across samples and thereby identification of differentially expressed genes. It is used in many biological contexts to characterize differences between cell lines and tissues, identify drug mode of action or drivers of drug resistance, among others. Genes can also be differentially regulated because of confounding factors that were not accounted for in the experimental plan, such as change in cell proliferation. Here, we identified genes for which expression consistently correlates with cell proliferation rates in proteomics and transcriptomics high-throughput data sets to determine the overall impact of cell growth rate on these data. We combined the analysis of 449 cell lines and 1,040 cell lines in five proteomics and three transcriptomics data sets to generate a refined list of 223 confounding genes that correlate with cell proliferation rates. These include many actors in DNA replication and mitosis, and genes periodically expressed during the cell cycle. It constitutes a valuable resource when analyzing high-throughput datasets showing changes in proliferation across conditions. We show how to use this resource to analyze *in vitro* drug screens and tumor samples. By disregarding the proliferation confounders, one can instead focus on the experiment-specific regulation events otherwise buried in the statistical analysis.

## INTRODUCTION

Nowadays, high-throughput proteome profiling allows relative quantification of thousands of proteins across samples. It is used in many biological contexts to characterize differences between cell lines and tissues, determine drug mode of actions, identify drivers of drug resistance, to name a few. While this reveals meaningful gene regulations across numerous conditions, these results can be confounded by secondary effects of a given treatment (or biological context). For example, a change in cell proliferation is a common undesired side effect of biological treatment and a well acknowledged confounding factor that influences results without being the intended effect of a given treatment^1^. Indeed, differences in cell growth rates correlate with the proportion of cells in each phase of the cell cycle: less proliferative cells have longer G1 or G2 phases than more proliferative cells. Consequently, slower-growing cell cultures will have more cells in G1 and G2 phase and fewer in S and M phase^2^, and S and M phase-specific proteins will thus be less abundant in the lysates.

Genes highly expressed in proliferative cells have been used as proliferation markers by pathologists and researchers for many years^3-5^. Their expression indeed often correlates with the proportion of cells in S and M phase in a given sample and can strongly correlate with tumor progression and prognosis^6^. Nevertheless, there is to our knowledge no study that determines which proteins confound hypothesis-free high-throughput data analysis by correlating with cell proliferation, and the overall impact of cell growth rate on the transcriptome and the proteome remains to be determined.

In this work, we first define a pseudo-proliferation index based on transcriptomics and proteomics data for cells with known proliferation rate. We use this to analyze even larger datasets to identify a list of genes that correlate with cell proliferation at both transcript and protein level. Like the Contaminant Repository for Affinity Purification (CRAPome)^7^, which is often used to highlight potential contaminant of pull-down mass spectrometry (MS) analyses, the list of confounding genes that we define here constitutes a valuable resource when analyzing datasets where proliferation could be affected. We illustrate this in the context of proteomics cancer classification and drug screens^6,8^, where identifying confounders of cell proliferation allows to quickly discard less relevant genes and focus on genes that are regulated in a more context-specific manner.

## RESULTS AND DISCUSSION

### Pseudo-proliferation index derived from transcriptomics and proteomics data

Cell doubling times, or proliferation rates, are rarely provided alongside proteomics and transcriptomics data, so calculating correlation between gene relative quantities and cell doubling times is only possible for a limited number of publicly available data sets. For this reason, we defined a list of proliferation markers which relative abundances reflect relative cell proliferation at protein and transcript level. The NCI60 cell lines^9^ have been extensively characterized with high-throughput proteomics^10-13^ and transcriptomics^14-16^ and their doubling times are publicly available (dtp.cancer.gov/discovery_development/nci-60/cell_list.htm; update of the 05/08/15). We used these data to identify proliferation markers that would reproducibly correlate with the inverse of doubling times (*i*.*e*. correlate with relative cell proliferation) in proteomes and transcriptomes.

We calculated the Pearson correlation to inverse doubling times for each of the 3,637 protein groups quantified in minimum three of the four NCI60 proteome data sets. Among these, we found seventeen human proteins that were reported as proliferation markers in the literature^3,17^. Most of these are transcribed at specific phases of the cell cycle^18^ (green line in Fig. 1a, and colored in Supplementary Figure S1). Although not referenced as cycling in Cyclebase v3.0, MCM3, MCM7 and MYBL2 have been shown to be expressed in a cell cycle-dependent fashion in single-cell transcriptomics^19^ where MCM3/7 and MYBL2 expression peaks in G1 and G2, respectively. In the same study, CCND1 is found cyclic at protein but not transcript level, peaking in G1.

**Figure 1:**
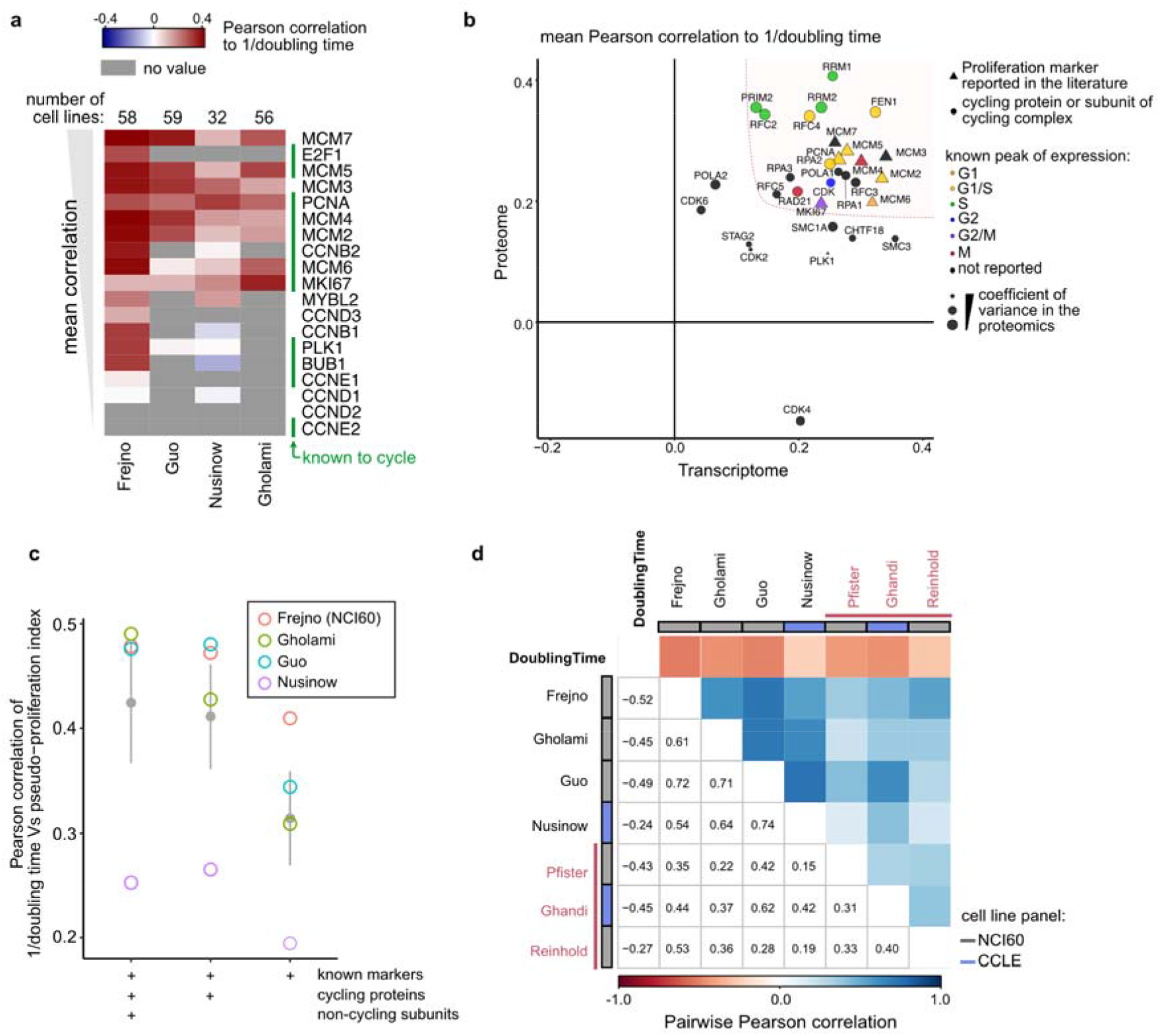
Calculation of pseudo-proliferation index. **a)** Pearson correlation with inverse doubling times of the NCI60 cell lines for proliferation markers referenced in the literature. The number of cell lines with known doubling times is indicated for each data set above the heatmap. The proteins that cycle in Cyclebase 3.0 are indicated by green bars on the right (“known to cycle”), and the data set names are indicated in the bottom. **b)** Mean Pearson correlation to inverse doubling time in the proteome (*y*-axis) and transcriptome (*x-*axis). The point size is proportional to the inverse of the coefficient of variance in the proteomics data, proteins present in less than 3 proteomics data sets were excluded. Periodic genes are color-coded by the phase of their expression peak, proliferation markers reported in the literature are indicated by triangles. **c)** Pearson correlations between pseudo-proliferation index and inverse doubling time in the proteomics data sets containing NCI60 cells using three sets of proliferation markers. Grey points and bars are mean and confidence intervals across data sets. **d)** Pairwise Pearson correlation between the pseudo-proliferation indexes calculated in the different data sets (proteomics and transcriptomics in black and red, respectively) and the doubling times provided for the NCI60 cell lines. Non-NCI60 cells were excluded from the data sets.

Figure 1a shows that the expression of most of these proliferation markers correlate strongly with the inverse of NCI60 cell doubling times. We hypothesized that other cycling genes could be good markers of cell proliferation, and that increasing the number of genes used to estimate cell proliferation would be more robust to missing values and quantification uncertainties. Among the genes known to cycle at transcript level according to ^18,20^, sixteen were quantified in minimum three of the NCI60 proteomics data sets (colored points in Supplementary Figure S1a). These proteins form complexes with other subunits that were not identified as cycling at RNA level but could correlate with the inverse doubling times; examples include the DNA polymerases A complex known to bind the cycling primases PRIM1 and PRIM2, or members of the replication factor C (RFC5 was not detected in ^20^ so its cycling status is unknown) (empty circles in Supplementary Fig. S1a).

Since we wanted to estimate relative cell proliferation in transcriptomics as well as in proteomics data, we also analyzed two transcriptomics data sets of the NCI60 cell lines^15,16^ (Supplementary Figure S1b). Figure 1b shows the correlation of the selected genes with inverse doubling time in the transcriptome (*x-*axis) and the proteome (*y*-axis). The periodic genes with the strongest correlation peak in G1/S and S phase at transcript and protein level, respectively^18^. From these data, we defined a preliminary set of potential proliferation markers containing the genes presenting high correlation with inverse doubling time both in the transcriptomics and proteomics data sets (Fig 1b, red area in the top-right corner). We compared pseudo-proliferation indexes calculated as the mean signal of:

- proliferation markers referenced in the literature (PCNA, MCM2–7 and MKI67).
- proliferation markers referenced in the literature and genes known to cycle at transcript level (FEN1, RRM1, RRM2, RAD21, CDK1, RPA2, RFC4, RFC2, PRIM2).
- all the above plus known interacting non-cycling subunits POLA1, RFC3, RPA1, RPA3, RFC5.

We compared how the resulting pseudo-proliferation indexes correlated with inverse doubling time in the proteomics NCI60 data sets (Fig. 1c). As expected, the more genes were included in the proliferation markers list, the stronger the correlation (except for the Nusinow et al. data set, which only contained 32 NCI60 cell lines). Based on these results, we decided to include the proliferation markers, periodic genes, and subunits of cycling complexes to calculate pseudo-proliferation index (all proteins in the top-right corner of Fig. 1b).

This data-driven approach was used to estimate relative cell proliferation on proteomics data sets with no doubling time reported: the proteomes of the CRC65 cancer cell lines^10^; and the Cancer Cell Line Encyclopedia (CCLE) that comprises the CRC65, NCI60 and other cell lines^13,21^. For each data set, we first calculated the pseudo-proliferation indexes, and next the correlations of each protein to this proxy for cell proliferation. Gene set enrichment analysis (GSEA) showed that proteins involved in chromatin remodeling, DNA replication and chromosome organization were highly correlated to pseudo-proliferation index (Supplementary Figure S2). These results were similar to those of a GSEA performed on proteins ranked by their correlation to inverse doubling time when available (“NCI60 only”), which confirmed that pseudo-proliferation index reflects the proliferative state of cells and can be used as an estimation of relative cell growth rates. We further controlled that the same Gene Ontology (GO) terms were reproducibly enriched across larger data sets of non-NCI60 cell lines. It is worth mentioning that the doubling times available on the NCI60 panel may be different from the actual doubling times in the different data sets analyzed here. Indeed, differences in experimental conditions and cell passages may affect cell growth rates^22^. This could explain why we observed lower *p*-values and higher normalized enrichment scores (NES) in the GSEA performed with pseudo-proliferation index than with inverse doubling times. With the same approach, we calculated pseudo-proliferation index at RNA level in data sets containing the NCI60 and CCLE cells transcriptome^14-16^ (Supplementary Table S2). Pseudo-proliferation indexes were highly consistent across proteomes (0.54 to 0.74) as well as between proteomes and transcriptomes while negatively correlating with the NCI60 doubling times (Figure 1d).

### Identification of proliferation confounders

Using pseudo-proliferation index, we could identify which protein quantities correlated with cell proliferation rates in the four proteomics data sets presented above (Supplementary Fig. S3a). We filtered out the proteins that were detected in less than two data sets and calculated the mean of Pearson correlations to pseudo-proliferation index across data sets.

We benchmarked our approach with three sets of genes expected to be confounders of cell proliferation (*i*.*e*. gold standards) either because they are known to be expressed in a cell-cycle-dependent fashion or because they were reported to be expressed under the control of a transcription factor only active on S-phase entry:

- B1: 48 genes known to be periodically expressed in synchronized cell cultures^23^.
- B2: 382 genes compiled from two list of proposed E2F transcription factor targets^20,24,25^.
- Cyclebase 3.0: 570 periodically-expressed genes (https://cyclebase.org/)^18^.

We ranked the proteins (excluding the proliferation markers used to calculate pseudo-proliferation index in the first place) by decreasing absolute mean of correlation to pseudo-proliferation index and counted the number of proteins belonging to each of the three gold standard sets (Fig. 2a). As expected, these gold standards were enriched in the top-ranked confounders (left of the *x-*axis), which indicates high absolute correlation with pseudo-proliferation index. We determined two cutoffs for low- and high-confidence correlation with pseudo-proliferation index: ≥ 0.313 and ≥ 0.385, respectively (“o” and “x” in Fig. 2a). The exact same strategy was applied with three transcriptomes to determine the transcriptomics confidence thresholds: ≥ 0.560 and ≥ 0.625 for low- and high-confidence, respectively (Fig. 2b). In both analyses, we calculated gene correlations with randomized pseudo-proliferation index (50 iterations) to check that all the low- and high-confidence cutoffs where under 0.1% FDR (see material and methods).

**Figure 2:**
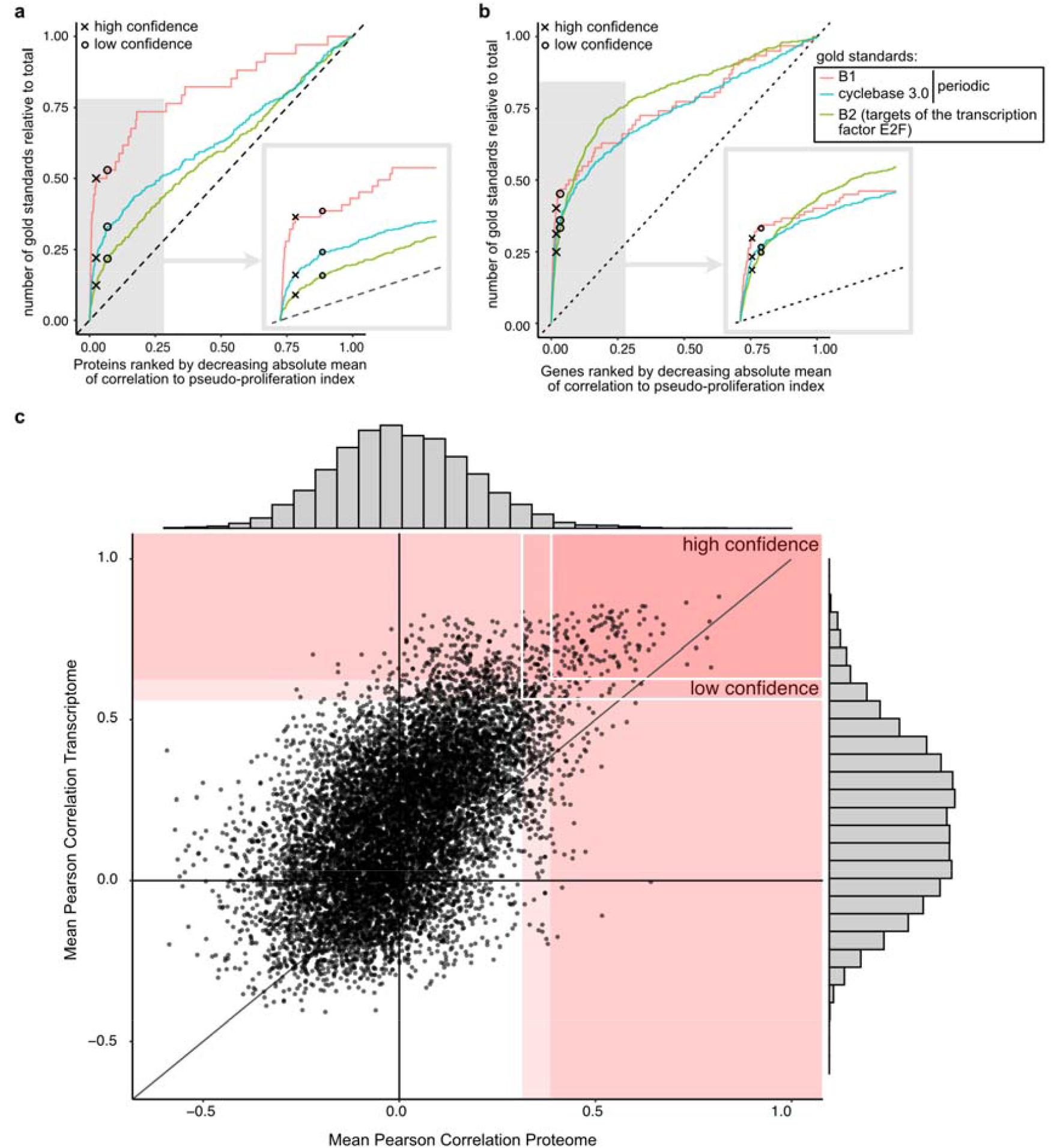
Confounding factors of cell proliferation. **a-b)** Definition of high- and low-confidence cutoffs for correlation with pseudo-proliferation index with three sets of gold standards in the proteomes (a) and the transcriptomes (b). Proteins/genes were ranked by decreasing absolute Pearson correlation to pseudo-proliferation index (*x*-axis) and the *y-*axis presents the cumulative number of gold standards for each set. Proteins/genes quantified in less than 3 and 2 data sets were excluded in (a) and (b), respectively. **c)** Scatter plot of the mean Pearson correlation to pseudo-proliferation index at protein (*x*-axis) and transcript (*y*-axis) level across all data sets. The red areas contain the proteins above the medium- and low-confidence thresholds in the proteome and/or transcriptome and the rectangles with white borders indicate the final list of low- and high-confidence proliferation confounders defined in this study. The point distribution in the proteomes and transcriptomes are presented on the sides of the plot.

Figures 2c shows gene correlations to pseudo-proliferation index at transcript and protein level. Overall, transcripts presented a higher mean correlation with pseudo-proliferation index than the proteins these were translated to, and the distribution of Pearson correlations to pseudo-proliferation index was wider at transcript than protein level. This indicates post-translational adjustment of protein quantities. Gene correlations to pseudo-proliferation index at protein and transcript level are available in Supplementary Table S3. We defined two set of confounding genes that correlate with pseudo-proliferation index at transcript as well as protein level (*i*.*e*. proliferation confounders): 223 and 119 genes were above the low- and high-confidence cutoffs in both proteomics and transcriptomics analysis, respectively.

Figure 3 shows the physical interactions between the high-confidence proliferation confounders identified in this study according to the STRING physical interaction subnetwork^9^. Each node (gene) is colored with its Pearson correlation with pseudo-proliferation index in each data set (ring). These confounding genes are involved in DNA replication and mitosis. As expected, we find back the genes used for calculating the pseudo-proliferation index (circled in black). Some of the proliferation markers previously described in the literature were high-confidence confounders, such as BUB1 and CCNB1/2 and PLK1 (dashed black borders in Fig. 3). These genes were not included in the refined list of proliferation markers used for calculating pseudo-proliferation index because they did not consistently correlate with the inverse of NCI60 doubling times, but they strongly correlate with relative cell proliferation when integrating more cell lines.

**Figure 3.**
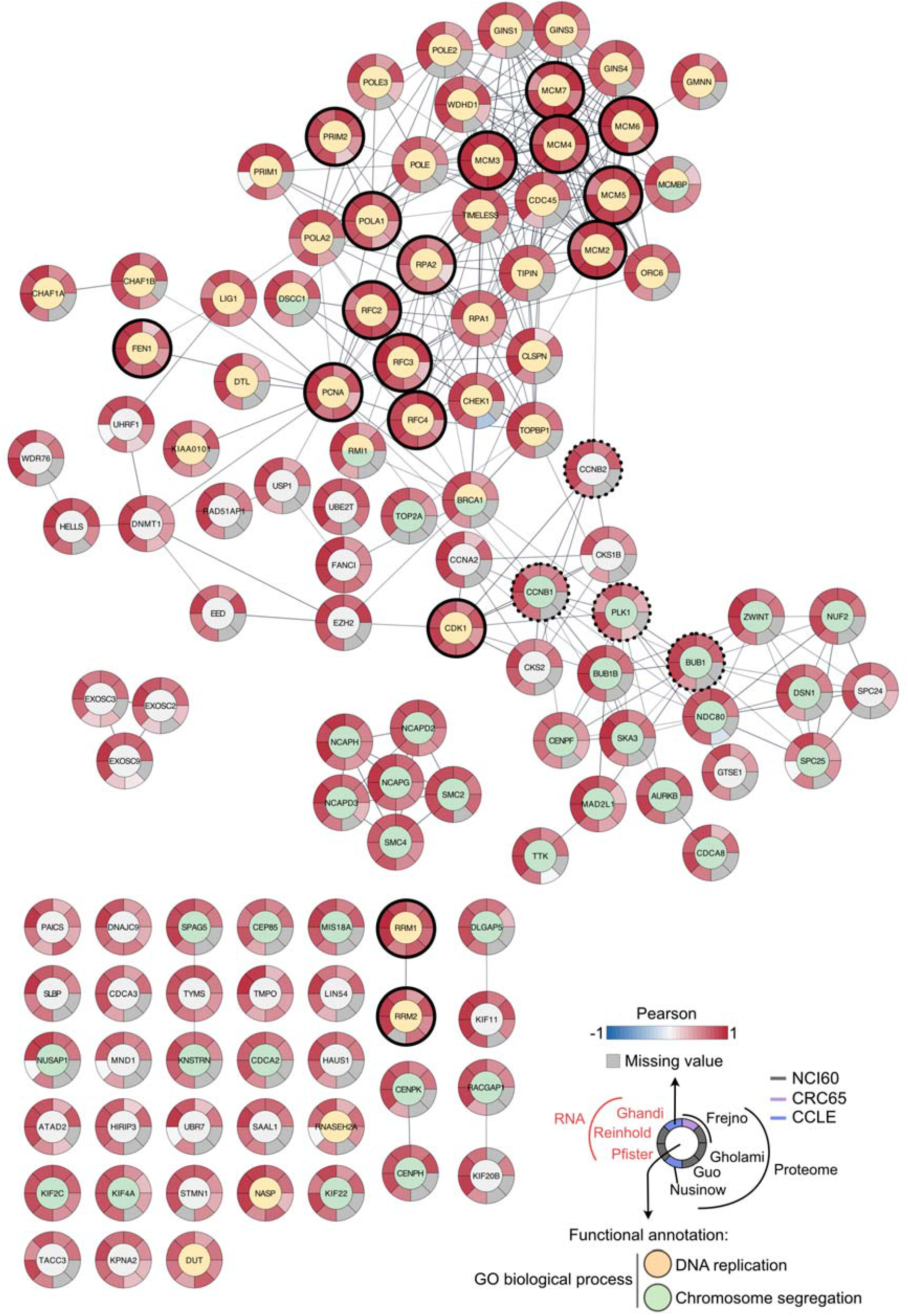
High-confidence proliferation confounders. STRING subnetwork of physical interactions (score ≥ 0.7) corresponding to the high-confidence proliferation confounders as selected in Fig. 2. The genes used to calculate the pseudo-proliferation index and the known proliferation markers not included for pseudo-proliferation index calculation are highlighted by black solid and dashed borders, respectively. The nodes are color-coded by selected gene annotations of biological processes. External ring are the Pearson correlations for each data set independently.

### Use case 1: Proliferation confounders in drug screens

Many drugs affect cell proliferation, thereby decreasing the proportion of cells actively dividing in samples. This can confuse data analysis when investigating drug mode of action because many of the genes regulated upon treatment are in fact confounding genes correlated with cell proliferation. A recently published paper provides the proteomes of five cell lines after 53 drug treatments^8^. In many experiments, proliferation confounders were enriched among the proteins that were downregulated after treatment, suggesting that the drug treatments reduced cell proliferation rates.

After befeldin A^26^ treatment, proliferation confounders are enriched among the most downregulated proteins (Fig. 4a). Brefeldin A disassembles the Golgi complex and induces endoplasmic reticulum (ER) stress. It is usually used as potent inhibitor of cell secretion. Consequently, Brefeldin A treatment reduces cell proliferation, which is very visible when labelling proliferation confounders in the volcano plot Figure 4b (red and orange dots): most of the cofounders are shifted towards the left of the volcano. Labelling them facilitates data analysis by: 1) highlighting global fold-change shifts that can be due to proliferation increase or decrease as a consequence of drug treatment and 2) disregarding protein regulations due to proliferation changes to concentrate on more direct consequences of drug treatment.

**Figure 4:**
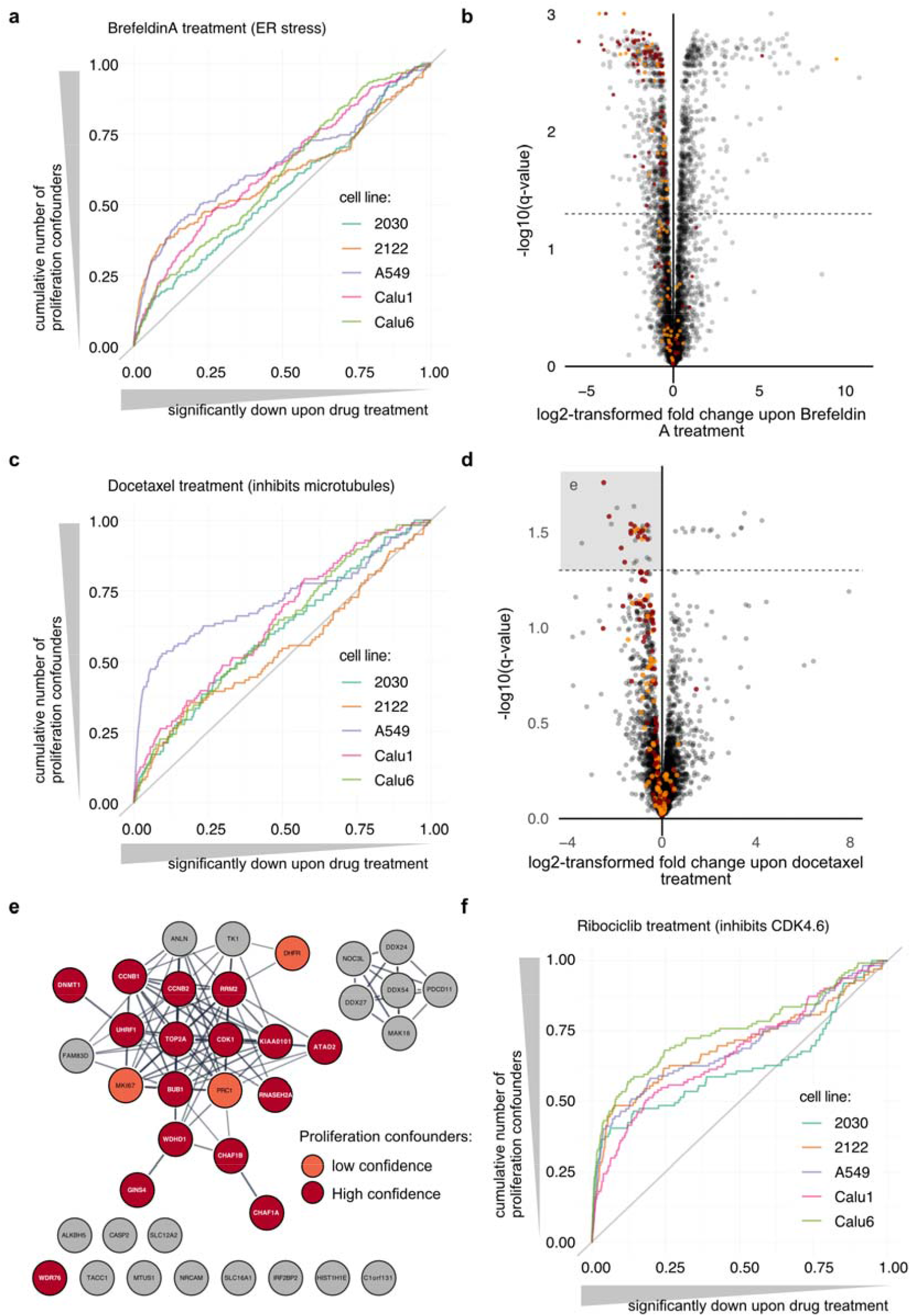
Proliferation confounders in the context of drug treatment. **a)** Enrichment of high-confidence proliferation confounders in the proteomes of cells treated with Brefeldin A. Proteins were ranked by significance of down-regulation according to Ruprecht et al. (q-value) (x-axis) and the y-axis presents the cumulative number of high-confidence confounder for each cell line. “2030” and “2122” correspond to the NCIH-2030 and NCIH-2122 cell lines, respectively. **b)** Volcano plot for A549 cells treated with Brefeldin A. High- and low-confidence proliferation confounders are highlighted in red and orange, respectively. The dashed line corresponds to a q-value of 0.05. **c)** Enrichment of high-confidence proliferation confounders in the proteomes of cells treated with Docotaxel as in (a). **d)** Volcano plot for A549 cells treated with Docetaxel as presented in (b). **e)** Significantly down-regulated proteins (grey square in (d)) are presented in a STRING network of functional associations (score ≥ 0.7). High- and low-confidence proliferation confounders are highlighted in red and orange, respectively. **f)** Enrichment of high-confidence proliferation confounders in the proteomes of cells treated with Ribociclib as in (a).

Docetaxel treatment impacts cell proliferation specifically in A549 cells (lung carcinoma epithelial cells) where downregulated proteins were enriched in high-confidence proliferation confounders (Fig. 4c). The volcano plot corresponding to this experiment is presented Figure 4d Docetaxel is a taxane that interferes with microtubule growth by binding to the β-subunit of tubulin. It is used in the treatment of many cancers. Here, we see that most of the proteins significantly downregulated upon treatment in A549 were proliferation confounders. Figure 4e shows the STRING network of functional associations of the proteins significantly downregulated in Figure 4d (grey box). Most of these genes are functionally connected in a “hairball” that contains all but one proliferation confounder. Some of these genes are involved in microtubule remodeling, but others are downregulated because of a reduction of cell proliferation of the A549 cells upon treatment. Examples of the latter include RRM2, which catalyzes the biosynthesis of deoxyribonucleotides, and the chromatin-assembly factors CHAF1A and CHAF1B. Labeling of the proliferation confounders facilitates the identification of proteins potentially more relevant to the drug treatment (grey nodes outside of the hairball). For example, the Microtubule-associated tumor suppressor 1 (ATIP3, coded by the gene *MTUS1*). *MUTS1-*deficiency is associated with increased microtubule dynamics^27^, which is the opposite of docetaxel-induced microtubule stabilization. In breast cancer, ATIP3 was found significantly downregulated in taxane-sensitive tumors^28^. It is an interesting therapeutic target for breast cancer^29^. Caspase 2 (*CASP2*) has been shown to cleave the Microtubule-associated protein tau (coded by the gene *MAPT*) that promotes microtubule assembly and stability and potentially competes with taxanes for microtubule binding. It is associated with resistance to taxanes in several cancers^30-32^. The Transforming acidic coiled-coil-containing protein 1 (TACC1) is also involved in microtubule regulation^33^. The Nucleolar complex protein 3 homolog (*NOC3L*), protein MAK16 homolog, RRP5 homolog (*PDCD11*) and the ATP-dependent RNA helicases DDX24/27/54 are RNA-binding proteins. Although there is no obvious known association of these proteins with docetaxel treatment and/or microtubule regulation, these downregulated proteins may inform on docetaxel impact on A549 cells.

In other cases, such as ribociclib treatment the same genes are not confounders but reflect the drug mode of action. Ribociclib inhibits CDK4/6 activity and thereby prevents progression through the G1/S checkpoint, blocking cells in G1 phase. This results in a high enrichment of proliferation confounders in negatively regulated genes (Fig. 4f and Supplementary Figure S3c), which is highly relevant for data interpretation.

### Use case 2: Proliferation confounders in the context of cancer prognostic and classification

The refined list of proliferation confounders can also be useful for analysis of *in vivo* samples and patient data, for example in the context of cancer since most tumors are characterized by an increased proliferation rate. Many of the confounding genes reported here are indeed reported prognostic markers in the context of cancer. Although this can be highly relevant, these proliferation confounders may only be significantly regulated because of the presence of more dividing cells in certain tumor samples. Other genes/proteins may be more appropriate for targeted therapy.

The recently published meta-analysis of the Clinical Proteomic Tumor Analysis Consortium (CPTAC)^6^ identified proteins which relative quantities are correlated with tumor grade or stage in patient samples. The high-confidence proliferation confounders identified in this study were not enriched in proteins associated with tumor stage (Fig. 5a, see Supplementary Figure S3d for lower confidence confounders). Proteins strongly correlating with tumor grade, however, were enriched in proliferation confounders in lung adenocarcinoma (LUAD), uterine endometrial carcinoma (UCEC), and pediatric glioma, but not in clear cell renal cell carcinoma (CCRCC) and ovarian serous adenocarcinoma (OV) (Fig. 5b, see Supplementary Figure S3e for lower confidence confounders). This is in agreement with the GO-term enrichment presented in Monsivais *et al*., where “cell cycle process” and “DNA replication” are strongly enriched in the proteins the most associated with cancer grades in LUAD, glioma and UCEC.

**Figure 5:**
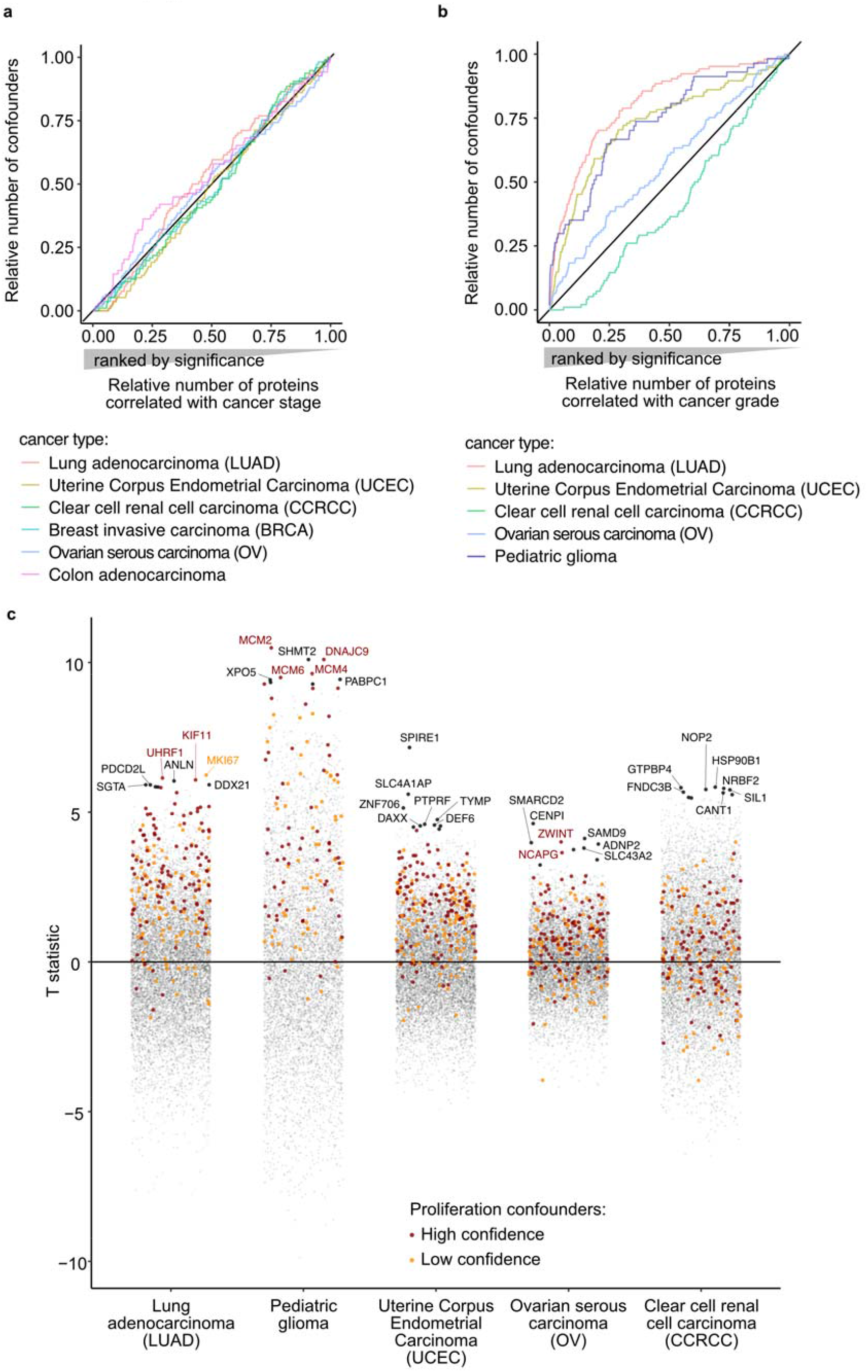
Proliferation confounders in the context of cancer grade. **a-b)** Enrichment of high-confidence proliferation confounders in proteins associated with cancer stage (a) and grade (b). Proteins were ranked by significance of correlation according to Monsivais et al. (*p*-value of Pearson correlation) (*x*-axis) and the *y-*axis presents the cumulative number of high-confidence confounder for each cancer type (see Supplementary Figure S3c-d for lower threshold). **d)** Proteins T-statistic provided by Monsivais et al. for the analysis of cancer grades (positive = high correlation with cancer grade) for each cancer type (*x-*axis). Each point corresponds to a protein, low- and high-confidence proliferation confounders are highlighted in orange and red, respectively. The seven top hits for each cancer type are indicated by their gene names.

In the lung adenocarcinoma and pediatric glioma data, the proteins the most associated with cancer grade include a high number of proliferation confounders that may not be the best candidates for targeted treatment. Figure 5c shows the proteins correlation with grade (*y-*axis), with high- and low-confidence proliferation confounders highlighted in red and orange, respectively. With such figure, it is possible to quickly identify proteins that are specifically correlated with high tumor grade but not associated with the high proliferative state of aggressive lung tumors.

In lung adenocarcinoma, the three proteins the most correlated to cancer grade were proliferation confounders: the U3 ubiquitin-protein ligase UHRF1, Kinesin-like protein KIF11 and the well-known proliferation marker MKI67 FHA domain-interacting nucleolar phosphoprotein. The Anillin actin binding protein (*ANLN*) was the top hit amongst non-confounding genes. It is highly expressed in lung cancer cell lines and tumor samples compared to normal tissues^34^, and *ANLN* high expression is a predictive marker of poor outcome for patients with lung adenocarcinoma in TCGA^35^. Anillin activates cellular migration of lung cancer cells *in vitro*^34^ and increases tumor growth and metastasis in breast cancer through induction of mesenchymal to epithelial trans-differentiation^36^. 67% of the high-confidence proliferation confounders identified in this study were correlated to cancer grade with a *p-*value under 0.01 in gliomas (Fig. 5c). In such context, it is particularly important to acknowledge that these genes may be regulated because of differences in cell growth rates. The confounders MCM2/4/6, and the heat shock co-chaperone and histone chaperone DNAJC9^37^ were amongst the five proteins the most correlated with glioma grades. In the figure, these surround the mitochondrial serine hydroxymethyltransferase (SHMT2) (ranked 3^rd^), which could be a more interesting hit for targeted therapy. It participates to the synthesis of glycine by catalyzing serine-to-glycine conversion. Glycine is a key resource for proliferative cells, and elevated concentration of glycine in IDH-mutated glioma tumors has been associated with aggressive glioma and is predictive of shortened patient survival^38^. While SHMT2 expression is not directly correlated with glycine concentration in gliomas^38,39^ it has been shown to favor cancer cells adaptation to poorly vascularized tumor micro-environments in the context of ischemic glioma^39^, and to be associated with poor prognostic in glioma^40^.

Highlighting proliferation confounders in the analysis of LUAD and pediatric glioma allowed to quickly focus on proteins more directly associated with cancer grades in these contexts. This illustrates the advantage of taking confounding genes into consideration when analyzing proteomics data of patient samples.

## CONCLUSION

Here, we calculated a pseudo-proliferation index that we used as proxy for relative cell proliferation at transcript and protein level to define low- and high-confidence thresholds for identifying confounding genes correlated with cell proliferation rates. We combined the transcriptomics and proteomics analysis to provide a final list of low- and high-confidence proliferation confounders. The Supplementary Table S3 provides the correlations to pseudo-proliferation index for 10,751 genes/proteins quantified in the data sets that were used for this analysis. With this list of proliferation confounders, anybody can identify in their data sets the genes/proteins correlated with cell proliferation like contaminants are routinely flagged using the CRAPome^7^.

We showed examples of high-throughput data analysis where labelling proliferation confounders facilitates data interpretation. We show that cell growth rate can be a confounding factor that results in down- or up-regulation of many genes in *in vitro* drug screens and tumor samples. Flagging these confounders among the most regulated genes allows to quickly identify other regulated hits that could be more relevant in the context of the experiment. Such analyses still require strong knowledge of the biological context and molecular regulation at play, but the genes correlated with cell proliferation rates are not all annotated as being involved in replication of cell-cycle-related processes. Thus, our refined list of proliferation confounders is an unvaluable resource for interpreting data where changes in cell growth rates/proliferation is a confounding factor.

The strategy that we describe here to identify proliferation confounders is straightforward and can be applied to many other types of confounding factors. The only requirement, which can be very limiting, is the availability of several high-dimensional data sets on samples where the confounding factor of interest can be quantified. We believe that taking confounding genes into consideration should become part of the high-throughput data analysis routine and will facilitate data interpretation in many biological contexts.

## MATERIALS AND METHODS

### Data availability

All the scripts and input tables associated to this study are available on Zenodo.org (10.5281/zenodo.6346643) under a BSD2-Clause “Simplified” license.

### Retrieval and pre-processing of proteomics data

The raw data from Gholami et al.^11^ were retrieved from the PRIDE proteomeXchange repository PXD005946 and searched against the Human reviewed protein database (download 12/03/2021 from Uniprot.org) with MaxQuant v1.6.17.0. The mqpar.xml and the fasta file associated with the search are available on Zenodo.org (10.5281/zenodo.6346643). The proteinGroups.txt table was filtered to remove the reverse sequences and potential contaminants identified with the contaminant database included in MaxQuant. We kept only the protein groups with minimum one unique peptide and a q-value ≤ 0.01 (6,900 protein groups). We further removed the samples with more than 70% of missing values. LFQ was utilized for correlation calculation after variance stabilizing normalization (vsn)^41^.

The data from Guo et al.^12^ were retrieved from the Table S1E provided in the paper (3,171 protein groups with no missing value) and normalized using vsn before correlation calculation.

The normalized iBAQ quantification from Frejno et al.^10^ was retrieved from the supplementary Data 3 available with the paper. The tables for Trypsin, GluC and Trypsin digestion of the CRC65 cells were filtered to remove the reverse sequences and potential contaminants identified with the contaminant database included in the original MaxQuant search. We kept only the protein groups with minimum one unique peptide and a q-value ≤ 0.01 (9,744 and 7,271 protein groups in the trypsin and GluC dataset for the NCi60 cells, respectively and 11,308 for the CRC65 digested with trypsin). We further removed the protein groups with more than 50% of missing values. We also removed the protein “PLIB” in the trypsin dataset due to bad annotation. For the analysis of the NCI60 cell lines, we took the protein groups mean signal from the trypsin and the GluC data sets.

The normalized TMT quantification from Nusinow et al.^13^ was retrieved from the supplementary data available on https://gygi.hms.harvard.edu/publications/ccle.html (“Protein Quantitation (TSV Format)”). The tables were filtered to remove the protein groups with more than 50% of missing values.

### Proteome inter-data set matching

Since the searches were performed on each proteomics data set independently, the same protein can be labelled differently in the search outputs (*i*.*e*. belong to different protein groups, split across several isoforms…). We retrieved the protein groups corresponding to the same protein in different data sets. We first combined variants/isoforms signal by keeping their mean values. Then, we matched and renamed them across data sets according to the mapping table that is provided as Supplementary Table S1. In the cases where several rows of a given data set were mapped to the same homogenized protein group ID, we kept the mean value per sample. If several accessions of a given data set corresponded to a unique accession in another data set, we favored the homogenized protein group ID with the highest number of matching protein groups across data sets. In cases of tie, we kept the one with the least “combined” accessions (several accessions corresponding to the homogenized accession in a given data set).

### Proteomics proliferation confounders

For each data set independently, we calculated the mean of signal of proteins of the MCM complex (MCM2, MCM7, MCM3, MCM4, MCM5 and MCM6), RAD21, CDK1, PCNA, RPA2, RRM1, RRM2, RFC4, RFC2, FEN1, MKI67, PRIM2, POLA1, RPA1, RPA3, RFC5, RFC3, to generate a pseudo-proliferation index. The gene names and corresponding Uniprot accessions are provide in Supplementary Table S4.

For each protein group quantified in a minimum of 10 cell lines, we calculated its Pearson correlation to cell lines pseudo-proliferation index and to 1/doubling time (available on dtp.cancer.gov/discovery_development/nci-60/cell_list.htm - Last Updated: 05/08/15). Missing values were replaced in each data set with the 1% quantile. We excluded the protein groups only quantified in one data set and calculated the mean of Pearson correlations. The absolute mean of Pearson correlation to pseudo-proliferation index was utilized to rank the protein groups. To identify high confidence proliferation confounders, we performed the same analysis after randomization of the cell lines’ pseudo-proliferation index (50 iterations). The distribution of the resulting absolute mean of Pearson correlations across data sets allowed us to define FDR thresholds: 0.1% FDR was obtained for an absolute mean of correlation to pseudo-proliferation index ≥ 0.239 in the proteomics data. To define high- and low-confidence thresholds, we benchmarked the list of proteins ranked by decreasing absolute Pearson correlation to pseudo-proliferation index with three gene lists of gold standards: B1^23^, B2^20,24,25^, and the periodic genes described in Cyclebase 3.0^18^. Their cumulative count in the ranked list of proteins was utilized to select Pearson correlation values corresponding to high enrichment of gold standards.

### Retrieval and processing of transcriptomics data

The processed data for the Affymetrix NCI60 dataset (Pfister et al.)^15^ was obtained from the GEO NCBI portal using the GeoQuery R package (v.2.60.0)^42^. Probesets of the HGU133Plus2 chip were mapped to Ensembl genes using the custom annotation provided by BrainArray^43^. The mapping file for probesets to ensembl genes was obtained from the BrainArray version 25 download page (brainarray.mbni.med.umich.edu). Probeset intensities were averaged across replicates of the same cell line. Only gene-specific probesets were considered: when multiple probesets corresponded to the same gene, the one with the highest mean signal across all cell lines was selected to represent the gene. In total, 16,554 Ensembl genes (of which 15,994 having the biotype “protein coding genes”) were uniquely mapped to probesets on the chip for the 59 NCI60 cell lines.

Raw fastq files corresponding to the NCI60 RNA-Seq profiling (Reinhold et al.)^16^ were obtained from the European Nucleotide Archive (project accession PRJNA433861). The raw sequence reads were trimmed using Trimmomatic v038^44^, using the adapter file “TruSeq3-PE-2.fa”, and with the following parameters: “LEADING:3 TRAILING:3 SLIDINGWINDOW:4:15 MINLEN:36”. Transcript abundance estimates were then obtained using salmon v1.4.0^45^ in “quant” mode with the default parameters against the Human GRCh38 cDNA set obtained from the Ensembl release 103^46^. Gene-level abundance estimates were summarized using the R package tximport v.1.20.0^47^, and upper quartile normalization was performed with the calcNormFactors function from the edgeR package v. 3.34.1^48^. Finally, expression levels were obtained for 57,937 Ensembl genes (21,391 having the biotype “protein coding genes”) for the same 59 cell lines profiled in Pfister *et al*..

For the CCLE dataset (Ghandi et al.)^14^, normalized gene expression levels in TPM (transcripts per million) units were obtained from the DepMap portal (depmap.org/portal, “CCLE_expression_full.csv”). We used the original Ensembl gene identifiers provided in the files: 51,832 Ensembl genes (19,790 protein coding) across 1,026 cell lines.

### Transcriptomics proliferation confounders

For each dataset independently, the pseudo-proliferation index was obtained as described for the proteomics datasets, by averaging the expression levels of the selected proliferation markers. For each gene, in each dataset we computed the correlations with the pseudo-proliferation index calculated for the dataset, as well as correlation with 1/doubling time using the NCI60 cell lines doubling times when available. We selected the genes quantified in at least two datasets and calculated the mean of Pearson correlations. We performed the same analysis after randomization of the cell lines’ pseudo-proliferation index (50 interations) to define FDR thresholds: 0.1% FDR was obtained for an absolute mean of correlation to pseudo-proliferation index ≥ 0.259 in the transcriptomics data.

Mapping between Ensembl gene identifiers and UniprotKB accessions has been performed using the UniProt.ws R package v2.32.0. For the integration of the RNA data with the proteome, we removed the genes from the transcriptome that matched to more than 6 protein groups in the proteome (9 genes) and reported the values of each gene from the transcriptome if they matched the same protein group (155 genes).

### Gene set enrichment and GO term redundancy reduction

Gene set enrichments were performed with R v4.0.3 (R-project.org/) and RStudio v1.3.1093 (rstudio.com/) on a x86_64-apple-darwin17.0 (64-bit) running macOS Big Sur 10.16, using the packages clusterProfiler v3.18.1^49^ and org.Hs.eg.db v 3.12.0. The protein accessions were ordered by decreasing Pearson correlation with 1/doubling time or proliferation index. We ran the function gseGO() with the following parameters: ont =“ALL”, keyType = “UNIPROT”, minGSSize = 6, maxGSSize = 800, pvalueCutoff = 0.05, verbose = TRUE, OrgDb = “org.Hs.eg.db”, eps = 0, pAdjustMethod = “BH”. The output summary was used to make the Supplementary Figure S2 that presents GSEA on data sets with only NCI60 cell lines (first 2 panels) or without any NCI60 cell (last panel). We then simplified the output to reduce GO terms redundancy globally: we calculated the pairwise Jaccard indexes between all pairs of GO terms identified across data sets. Pairs of GO terms with a Jaccard index ≥ 0.5 were considered similar and only the one with the lowest enrichment *q-* value in any data set was kept for plotting. Figure S2 shows the 80 biological processes with the lowest absolute *q-*value (minimum value across all data sets and enrichments).

### Functional annotations and networks

The two gene/protein networks presented in this paper were generated with Cytoscape v 3.9.1^50^. GO term annotations were retrieved with the StringApp v 1.7.0^51^ and the donut visualization of Pearson correlations was performed with Omics Visualizer v 1.3.0^52^.

### Proliferation confounders in the context of drug treatment

The proteomics analyses of drug-treated cells were found in the supplementary Data 1 of Ruprecht et al.^8^. We counted the cumulative number of proteins subjected to statistical analysis by the authors and with a negative fold change upon drug treatment with Ribociclib (10,000 nM in all cell lines), Brefeldin A (100 nM, 30 nM, 100 nM, 100 nM, 30 nM for NHI-2030, NHI-2122, A549, Calu1 and Calu6, respectively) and Docetaxel (30 nM, 3 nM, 1 nM, 10 nM, 3 nM for NHI-2030, NHI-2122, A549, Calu1 and Calu6, respectively). Volcano plots were drawn with the data from Ruprecht et al.^8^, proliferation confounders were mapped to protein groups if minimum one of the protein in the protein groups had a gene name corresponding to a confounder.

### Proliferation confounders in the context of cancer tissue samples

The proteomics analyses of tumor tissues were found in Montsivais et al.^6^ (Supplementary Data 2 and 3 for correlation with grade and stages, respectively). Proliferation confounders were mapped to protein groups if minimum one of the proteins in the protein groups had a gene name corresponding to a confounder.

## CONTRIBUTION OF THE AUTHORS

**Marie Locard-Paulet:** Conceptualization, Methodology, Formal analysis, Investigation, Writing - Original draft, Visualization. **Oana Palasca:** Conceptualization, Methodology, Formal analysis, Investigation, Writing - Review & editing. **Lars Juhl Jensen:** Conceptualization, Methodology, Writing - Review & editing, Supervision, Funding acquisition.

## ACKNOWLEDGEMENT

Novo Nordisk Foundation Center for Protein Research is supported financially by the Novo Nordisk Foundation (Grant agreement NNF14CC0001).

