## Supplementary Figures for "Identifying the genes impacted by cell proliferation in proteomics and transcriptomics studies"

### Supplementary figures associated to the paper: Identifying the genes impacted by cell proliferation in proteomics and transcriptomics studies

Marie Locard-Paulet[1], Oana Palasca[1], Lars Juhl Jensen[1].

[1] Novo Nordisk Foundation Center for Protein Research, University of Copenhagen, Denmark

Locard-Paulet et al., Supplementary Figure S1

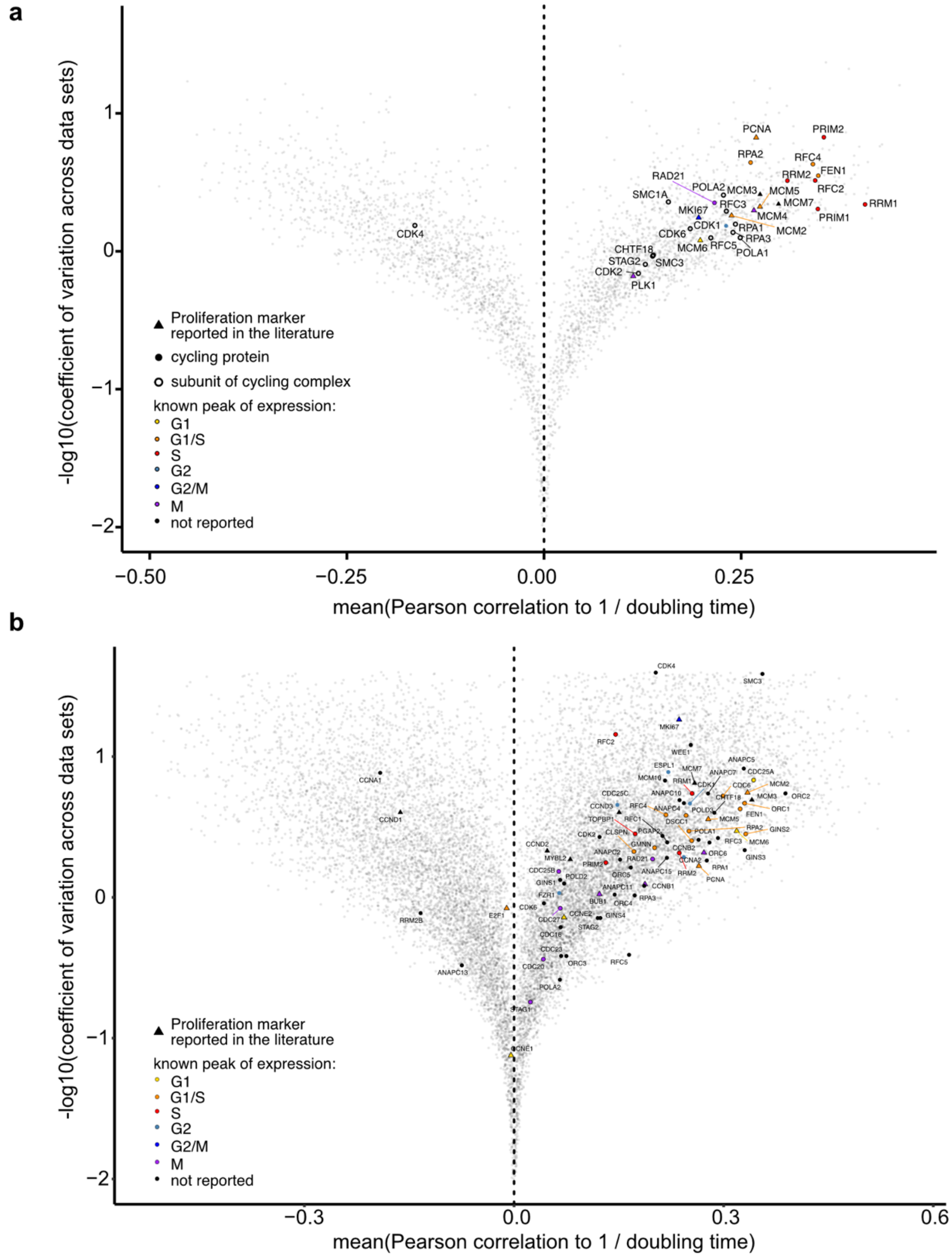


**Figure S1: Selection of proliferation markers for calculating pseudo-proliferation index in proteomics and transcriptomics data.** Volcano plots showing the mean correlation of proteins (a) or transcripts (b) to inverse doubling time in the NCI60 data sets (*x*-axis) and the -log10(coefficient of variance) across all the data sets (*y*-axis). Proteins quantified in less than 3 data sets were excluded in (a). Proteins/genes of interest are highlighted, and proliferation markers identified from literature search are indicated with triangles. These were color coded based of their expression peak according to (Santos, Wernersson et al. 2015).

Locard-Paulet et al., Supplementary Figure S2

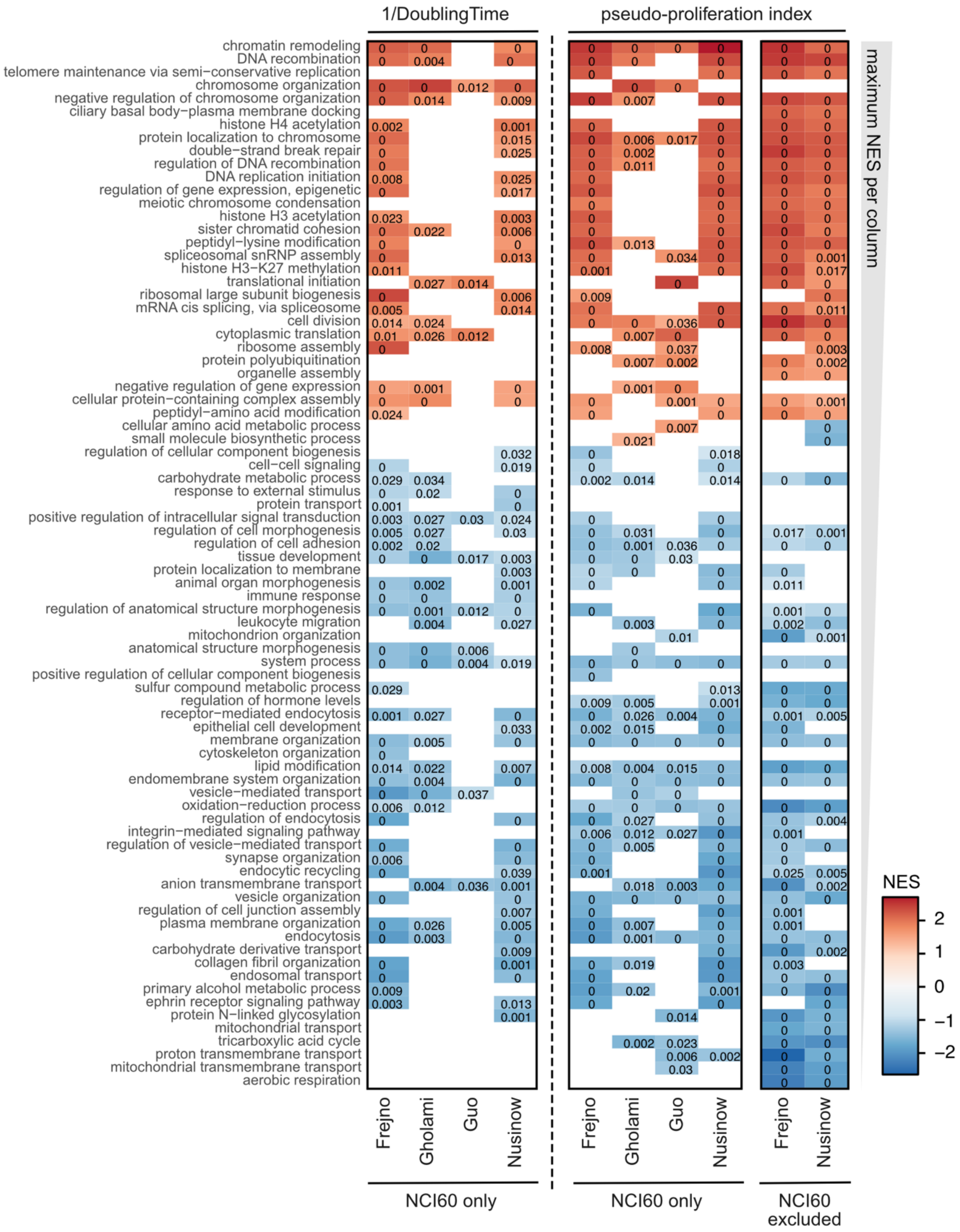


**<u>Figure S2:</u> Comparison of the biological functions enriched in proteins strongly correlated with inverse cell lines doubling times or pseudo-proliferation index.** For each data set, genes (with the exception of the genes used for calculating pseudo-proliferation index) were ranked based on their correlation to inverse doubling time or correlation to pseudo-proliferation index (left and right panel, respectively). Gene set enrichments were performed using the “gseGO” function from the R package clusterProfiler v 3.18.1, resulting *p*-values are indicated in each tile, as well as color-coded normalized enrichment scores

(NES). Only the annotations from biological processes are included, they are ordered by decreasing maximum NES per data set (top 80 enriched GO terms, see material and methods for a detailed description of the procedure used to reduce GO redundancy). Data sets are labeled based on the first author's name, enrichments were performed independently on the NCI60 cell lines or cell lines with no reported doubling time ("NCI60 only" and "NCI60 excluded", respectively).

Locard-Paulet et al., Supplementary Figure S3

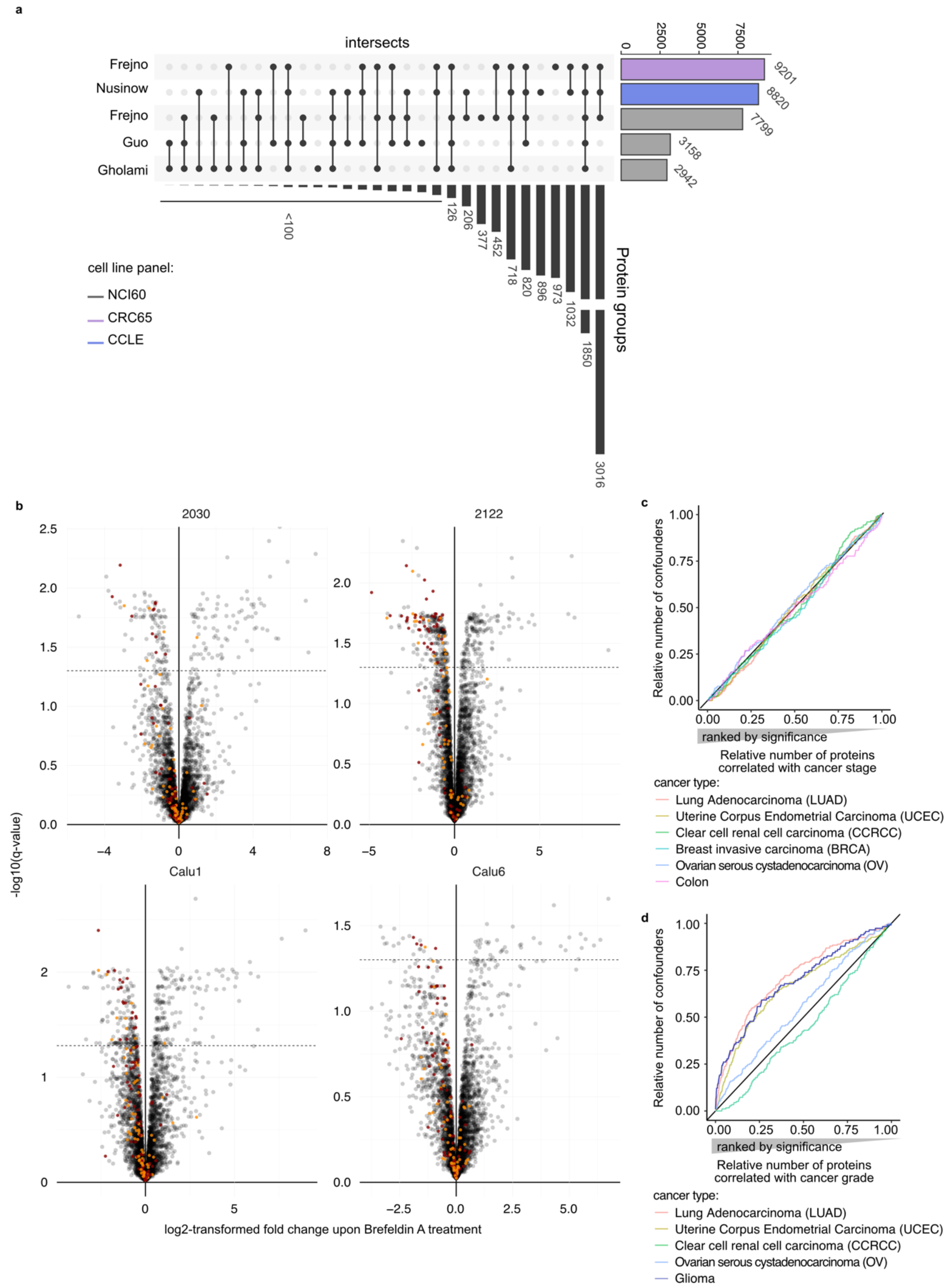


**Supplementary Figure S3. Proteomics data and use cases. a)** Protein coverage of the proteomics data sets used in the study (after isoform removal and accessions homogenization - see material and methods). These are identified by the first author’s name (bottom). The data set “Frejno” contained two independent MS searches of different cell line panels, we kept them separated. The total number of protein groups detected in the data sets are indicated in the top bar plot (color-coded by the cell line panel: purple, blue and grey for the CRC65, CCLE and NCI60 panel, respectively). The protein groups identified in multiple data sets are indicated by the bar plot on the right-hand side: number of protein groups detected in the data sets indicated by a dot on the dot plot. **b)** Volcano plots for each cell line treated with Brefeldin A. High- and low-confidence proliferation confounders are highlighted in red and orange, respectively. The dashed line corresponds to a *q*-value of 0.05. **c-d)** Enrichment of high- and low-confidence proliferation confounders in the proteins correlated with cancer stage (c) or grade (d) in Monsivais et al.. Lines are color-coded by cancer type. Axis are normalized to the total number of confounders and protein groups.
